# Control of spatio-temporal patterning via cell density in a multicellular synthetic gene circuit

**DOI:** 10.1101/2022.10.04.510900

**Authors:** Marco Santorelli, Pranav S. Bhamidipati, Andriu Kavanagh, Victoria A. MacKrell, Trusha Sondkar, Matt Thomson, Leonardo Morsut

**Author notes:** These authors contributed equally (Marco Santorelli, Pranav S. Bhamidipati).

## Abstract

A major goal in synthetic development is to design and construct gene regulatory circuits that control the patterning and morphogenesis of synthetic multicellular structures. In natural development, an interplay between mechanical and chemical communication shapes the dynamics of gene regulatory circuits that underlie patterning and morphogenesis. However, for synthetic gene circuits, how the non-genetic properties of the growth environment impact circuit behavior remains poorly understood. Here, we describe an occurrence of mechano-chemical coupling in synthetic contact-dependent synNotch patterning circuits demonstrating that cell density modulates the transduction of signal between a sender and receiver cell. By exploiting density-dependent signaling, we construct multicellular signal propagation circuits with synNotch and control the patterning outcome both temporally and spatially via cell density gradients established in vitro via plating or small-molecule mediated modulation of proliferation. Our work demonstrates that synthetic gene circuits can be critically impacted by their context, providing an alternate means for programming multi-cellular circuit patterning outcomes.

## Introduction

Morphogenesis emerges through the interplay between chemical and mechanical processes that occur simultaneously in development to generate the architecture of the embryo^1^. During development, chemical and mechanical processes can proceed in sequence with patterning providing a template for mechanical regulation. For example, in developing embryos, gene regulatory networks interact with intracellular signaling events to pattern domains of gene expression (e.g. Drosophila early development^2^). These domains can be characterized by the induction of motor proteins or differential adhesivity that deform cell shape and change embryonic geometry, driving morphogenetic events like gastrulation^3^ or germ-band extension^4,5^. However, recent work demonstrates that mechanical and chemical events can also be coupled in the other direction. Signaling pathways like the Yap/Taz axis can sense mechanical cues and convert these directly into changes in gene expression^6^. Similarly mechanical changes in the fluidity of a tissue can yield changes in gene expression dynamics^7–9^. Finally, the two aspects (mechanics and chemical) can be intertwined in so-called mechano-chemical systems, that seem to abound in developmental transitions^10–20^. Despite emerging examples, many principles remain obscure regarding how information flows between mechanical processes and chemical circuits in general, and how this contributes to expand, constrain or regulate patterning and morphogenetic outcomes.

The complexity of the embryo generates inherent challenges in studying and extracting general principles both of developmental transitions in general, and of interactions between mechanical properties and gene circuit signaling dynamics in particular. A major emerging theme in the field of synthetic development is the construction of gene circuits that enable controlled morphogenesis of synthetic embryos and organoids. Highly simplified engineered systems have been generated in this fashion that provide controlled and defined experimental systems in which to analyze gene circuits within the context of a multicellular structure^17,21–29^. Engineered systems provide an ideal setting to study mechano-chemical coupling where signaling and mechanical phenomena can be isolated, measured and modulated.

Synthetic circuits based on notch signaling, synNotch circuits, have emerged as a modular and flexible strategy for engineering multicellular mammalian systems^30–32^. The synNotch system uses engineered receptors modeled after the endogenous developmental signaling pathway Notch/Delta. This endogenous pathway is contact-dependent and is used extensively during development to generate cell-scale patterns. In the synthetic version, synNotch, both the input and output of the pathway have been rendered user-definable and, as such, are orthogonal to the endogenous Notch pathway. Using this system, developmental circuits have been developed in 2D in epithelial cells, and in 3D in fibroblast aggregates where a synthetic signal affects multicellular signaling and mechanics, for example, by driving expression of key adhesion proteins in the cadherin family [22, 30, 32]. By changing the nature of the adhesion proteins, or the circuits of cell-cell communication, a variety of different developmental trajectories have been implemented.

In the synNotch adhesion example, information flows from engineered synthetic signaling to changes in mechanical properties of the cellular system through changes in cell-cell adhesion. To achieve a complete synthetic mechano-chemical signaling system with reciprocal information flow between mechanical and chemical signals, mechanical inputs to signaling will have to be incorporated. Insights that synNotch could be a good candidate to develop such a system are emerging. First, the various proposed mechanisms of activation for Notch and synNotch signaling involve a mechanical “pulling force” that exposes the protease cleavage site for further signal transduction^33–37^, although the specific mechanisms may differ based on the cellular context and the endogenous or synthetic nature of the receptors^38^. Second, cellular mechanical tension, shear stress, and ECM stiffness have been shown to play a role in Notch signaling in certain contexts on the sender cell side or the receiver cell side^39–44^. Third, synNotch has been engineered to respond to different degrees of pulling force, highlighting the mechanosensory potential of this signaling modality^45^. Whether these inputs can be used to create a system where mechanics affects not only signaling, but also patterning outcomes has not been explored.

Mathematical models have been an important tool for understanding morphogenesis in natural systems^46–48^ and thus provide a potential strategy for the design and analysis of synthetic systems that incorporate mechanical-chemical coupling. Cell-based models of Notch-mediated signaling^49^ have uncovered key insights into the self-organization of regular spatial patterns^50^, the regulation of cell fate bifurcation by receptor-ligand interactions and cell geometry^51–53^, and the important roles of ligand expression levels and competition in robust patterning^54,55^. In addition to endogenous signaling, mathematical modeling of synthetic (exogenous) signaling networks has been a key tool for engineering exquisitely defined, controllable biological circuitry. Recently, cell-based modeling has been used similarly to catalyze the discovery and design of novel circuits for morphogenesis^56,57^. Such models have been used to study natural cases of mechano-chemical coupling^53,58–60^ but have not yet been applied in the synthetic context.

A deeper analysis of mechanical-chemical coupling in synthetic gene circuits could provide new insights into the logic of these circuits, and novel strategies for engineering multicellular system patterning and morphogenesis. Here, we first identify cell density as a non-genetic parameter that affects synNotch signaling, through a screening of mechanical inputs. We then apply the synNotch system to study the impact of cell density on a paradigmatic example of signaling-mediated patterning, i.e. lateral propagation via contact-dependent signaling. We construct multicellular tissues containing a local relay circuit consisting of signaling initiating, sender, cells and transceiver cells that both send and receive signals. Using this simplified genetic circuit, we investigate the impact of non-genetic inputs on patterning and signal propagation at the multicellular level. With a combination of in silico and in vitro experiments, we show that density modulates patterning in a parameter space that is biologically relevant and can be exploited to construct distinct macroscopic spatial and temporal patterns.

## Results

### Cell density impacts SynNotch signal transduction

The impact of non-genetic factors including tissue mechanics on synNotch signal propagation has not been explored. To quantify the impact of individual perturbations to the physical environment on synNotch signaling, we used a previously reported in vitro assay for synNotch activation based on a sender-receiver cell signaling paradigm. Briefly, two L929 mouse fibroblast cell lines, a sender cell line and a receiver cell line, are engineered such that contact between a sender cell and receiver cell can be assessed by the presence of a fluorescent reporter in receiver cells. Sender cells constitutively express membrane-bound green fluorescent protein (GFP), which acts as the ligand for an anti-GFP synNotch receptor on receiver cells (anti-GFP synNotch, Figure 1A). The intracellular portion of the anti-GFP synNotch receptor contains a tetracycline-controlled transactivator (tTA) which is freed from the membrane upon contact-dependent activation and translocates to the nucleus where it activates expression of cytosolic mCherry. To assay synNotch activity, sender and receiver fibroblasts are co-cultured in a 1:1 ratio for 24 hours, by which time activated receiver cells produce mCherry (Figure 1B). The control activation is performed on tissue-culture treated plastic dishes, at a confluent density where all cells are in contact with each-other, with no drugs. In the positive and negative controls (labeled “ON” and “OFF”) sender cells are present and absent, respectively. Expression of mCherry is quantified by fluorescence-activated cell sorting (FACS) at 24 hrs, and we compare activation levels in the control conditions to varied conditions of extracellular matrix (ECM) composition, substrate stiffness, cytoskeletal tension, and cell density, modulating each parameter individually. We report and refer to the density of the cell culture as a multiple of the density at 100% confluence (surface is 100% covered with cells). We estimated by visual inspection that the confluent density is 1250 cells/mm^2^ and refer to it as 1× confluence or simply “1x”.

**Figure 1:**
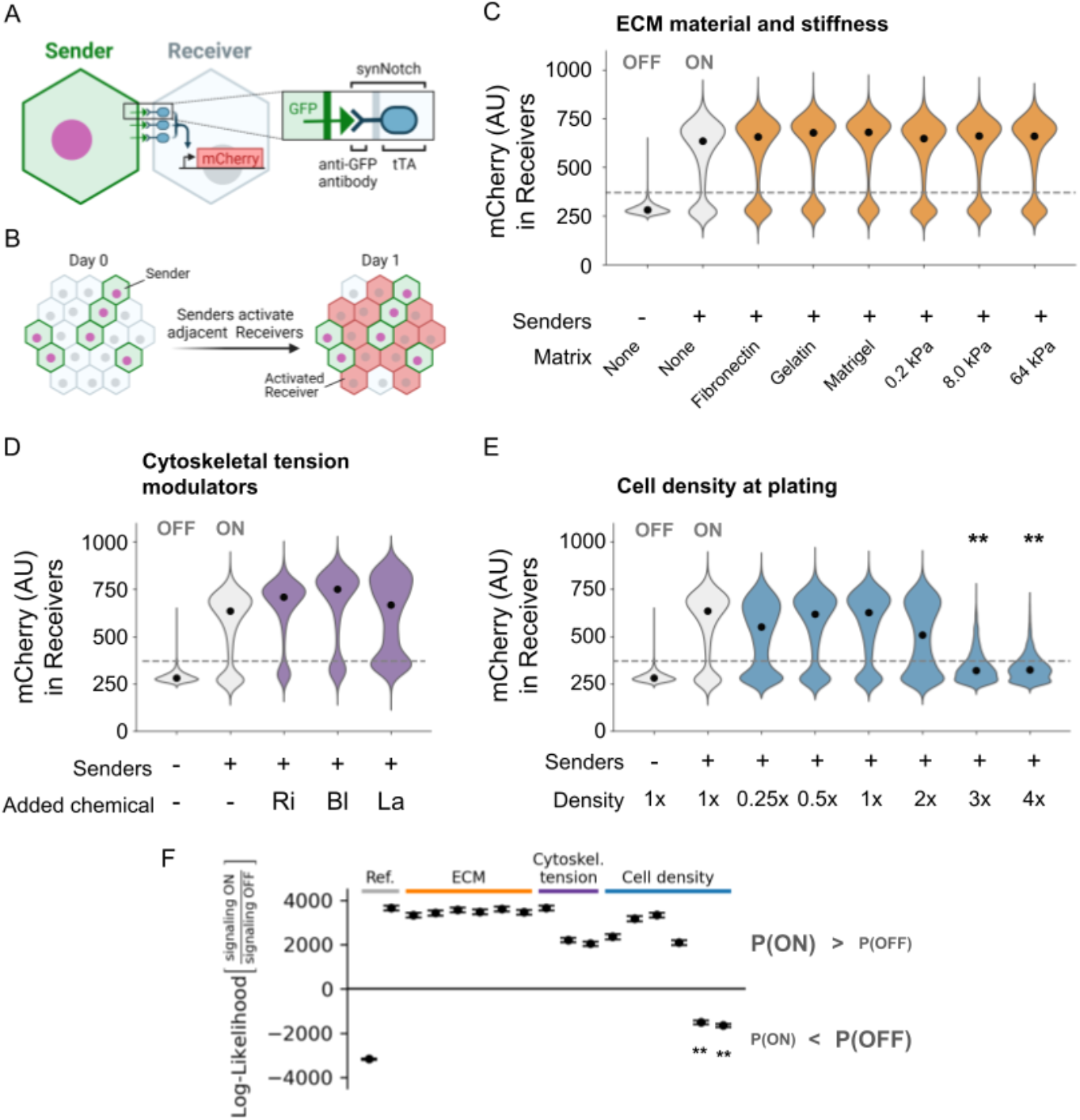
SynNotch activity depends on cell density, not ECM or cytoskeletal tension. (A) Conceptual scheme of Sender-Receiver synNotch signaling. Membrane-bound GFP (green triangle) in Sender cells binds synNotch in Receiver cells on the right. Cleavage of synNotch frees the intracellular domain (tTA, blue ellipsoid) to translocate to the nucleus and activate mCherry reporter expression. (B) Schematic depiction of synNotch signaling response assay. Senders and Receivers are co-cultured at 1:1 ratio, and synNotch signaling occurs at Sender-Receiver contacts (activated Receivers in red). Receiver activation (mCherry fluorescence) is measured at 24 hours by FACS under different culture conditions (C-E). Each violin in (C-E) depicts a distribution of mCherry fluorescence in Receivers. Gray violins are the reference samples for OFF and ON Receiver states (without and with Senders, respectively). Black dots indicate median. Dotted line indicates the cell-wise threshold between OFF and ON states. ** indicates the sample is more likely OFF than ON, as determined by the log-likelihood (LR) statistical comparison. Distributions are shown across (C) various growth substrate materials and stiffnesses, (D) various chemical modulators of cytoskeletal tension†, and (E) various initial cell densities. (F) Scatter plot of the LLR calculated for each sample. Points above zero indicate the sample resembles the ON state more than the OFF state. Error bars represent 95% confidence intervals (calculated by bootstrapping). For details please refer to Methods: Experiments. †RI: 100*µ*M ROCK-inhibitor (Y-27632); BL: 25*µ*g/mL blebbistatin; LA: 200*µ*M latrunculin-A.

We found that increased cell density, but not perturbations to other aspects of the physical cell environment, led to attenuation of synNotch signalling (Fig. 1C-F). Cells grown on different substrates, mimicking different ECM compositions, or at varying stiffnesses exhibited similar mCherry activation to reference receiver cells activated by senders (Fig. 1C, Methods: Experiments). Cytoskeletal tension was modulated by the addition of three drugs known to affect cytoskeletal contractility and actin polymerization (Y-27632 (ROCK-inhibitor)^61^, blebbistatin^62^, and latrunculin-A^63,64^). While these treatments may slightly increase synNotch signaling activity, they also increase basal activation in the absence of input ligand (Fig 1D and S1D). In contrast, when cells were grown across a range of cell densities from 0.25x to 4x confluence (where 1x is the density of a 100% confluent culture), higher densities resulted in a significantly inhibited signaling response in receivers (Fig. 1E), whereas lower densities showed slightly inhibited signaling activity. For details on experimental methods see Methods: Experiments. These results suggested that synNotch signal transduction is optimally efficient at cell densities of 0.5x - 1.0x (625 - 1250 cells/mm^2^), whereas much lower densities or much higher densities result in blunted signal transduction. For all treatment conditions, fluorescence was compared to histograms of unperturbed reference samples in the presence or absence of sender cells (labeled “ON” and “OFF”, respectively in Fig. 1C-E). A perturbation was considered inhibitory (labeled **) if its observed fluorescence values at 24 hours was more likely to arise from the OFF condition than the ON condition as measured by the likelihood ratio (LR), a parameter-free statistical comparison (Figure 1F). For details on calculations used, see Methods: Statistical analysis.

The observed decrease of synNotch signaling at cellular densities above 1x could be due to a number of molecular mechanisms. We investigated the nature of the underlying mechanisms by collecting observations of measurable variables that differ at 1x and 4x confluencies. As shown in Figure S3, we find that certain parameters change at density higher than 1x: (i) cells change shape as a function of density, becoming smaller surface-wise (Fig. S3A) and volume-wise (Fig. S3B), and surface curvature also increases at high density (Fig. S3C); (ii) membrane-targeted GFP ligand expression decreases sharply with density in a way that correlates with density-dependence of signaling (Fig. S3D-F); we find a positive correlation between GFP ligand expression and cell diameter (bigger cells, more ligand, smaller cells less ligand, Fig. S3G-H); (iii) cell motility decreases with increasing cell density (Fig. S3I). Other parameters do not change with confluency: (i) YAP does not change localization (Fig. S3J); (ii) conditioned media does not affect signaling (Fig. S3K); and (iii) nuclear reporter protein accumulation levels do not change (Fig. S3L). For further details, see Methods: Experiments.

Our measurements were most consistent with a mechanism in which density dependent signal attenuation occurs through modulation of membrane protein trafficking. Given that YAP localization does not change, and cytoskeletal drugs do not affect signaling, it seems unlikely that the decrease in signaling at high densities is due to classic cytoskeletal-tension-mediated mechanotransduction. Given that conditioned media from high-confluency cultures does not perturb signaling at low confluency, this does not seem to be due to soluble chemical factors. Given that the membrane reporter but not the nuclear reporter changed their apparent abundance over time, we hypothesize local, curvature-based changes in membrane trafficking, that results in higher protein turnover at higher curvatures, as observed for other signaling molecules^65–67^. Moreover, a role in cell motility can also be part of this phenomenon. Given that the nature of the mechanism seems to be on the membrane features, pointing to this being local/mechanical in nature and not chemical, we wondered if this phenomenon could be exploited to control outcomes of synNotch-mediated patterning in time and space.

### Signal propagation circuit exhibits density-dependency patterning outcome

We next sought to investigate the impact of density-dependent signal attenuation on the behavior of a multicellular synNotch patterning circuit. As one paradigmatic example of cell-cell contact dependent patterning, we chose to focus on the so-called “lateral propagation” circuit, where a signal is passed via contact-dependent signaling between neighboring cells. The lateral propagation circuit is a well-known developmental circuit^68^. It has also been implemented in a cellular system with a semi-synthetic system^69^, but the possibility to modulate its outcomes with changes in cell density has not been evaluated. The logic of the circuit is shown in Figure **2**A. The circuit relies on cells that can both receive and send a signal, or “Transceiver” cells. When in contact with one another and activated by a Sender cell, Transceiver cells become activated and are able to relay the signal to neighboring Transceivers in a propagating wave of signaling. An initial Sender cell is required to initiate signal propagation; however, thereafter Transceivers are capable of propagating the signal without additional stimulation.

**Figure 2:**
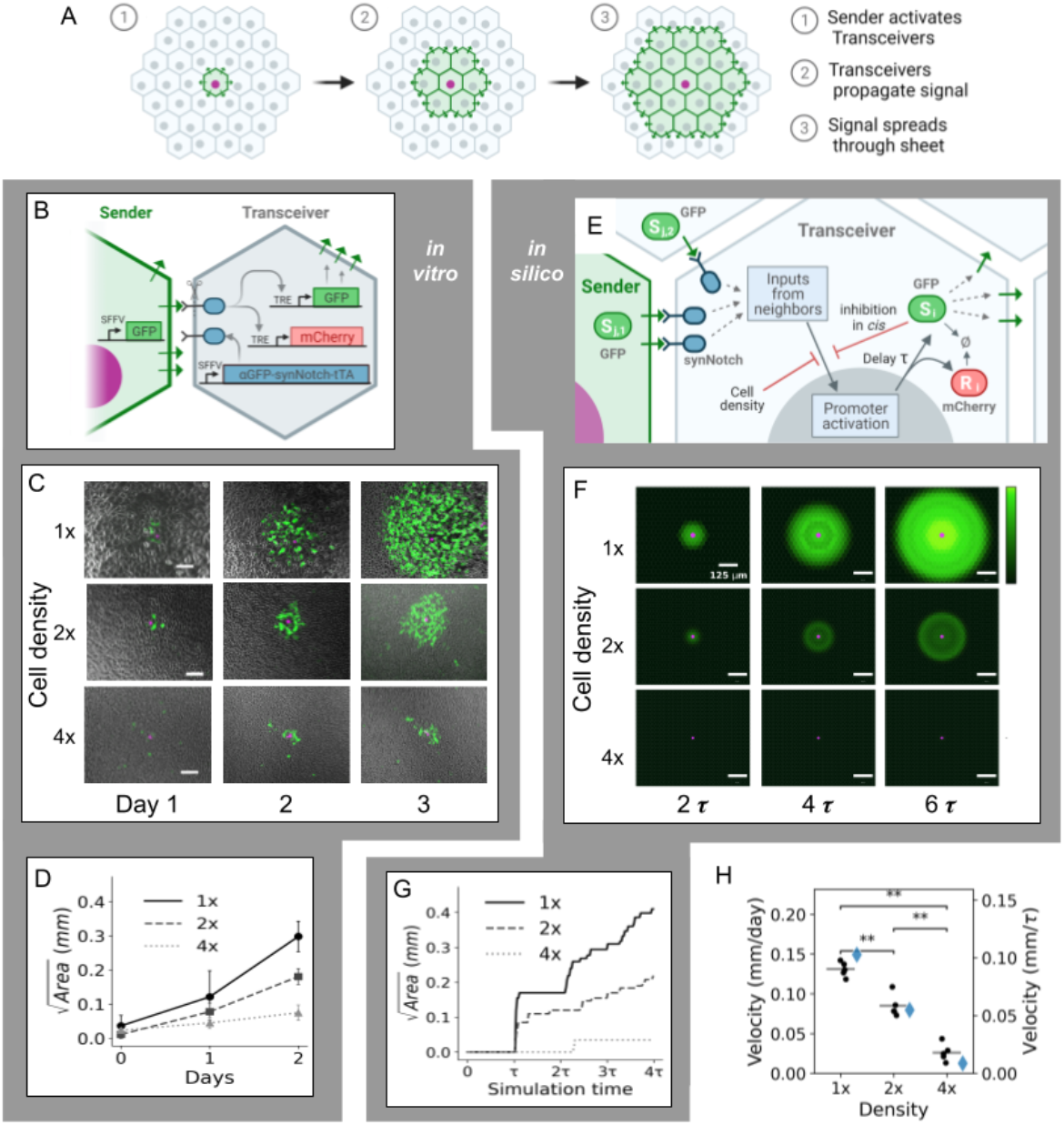
Cell density tunes the velocity of signal propagation. (A) Schematic of signaling wave propagation (green) in a monolayer of transceiver cells (gray) initiated by a sender cell (purple nucleus).(B) Schematic of the *in vitro* Transceiver circuit. The Sender cell (green) contains a transgene for constitutive expression of membrane-bound GFP ligand (green triangle) and a nuclear infrared fluorescent marker (not shown) under control of spleen focus-forming virus (SFFV) promoter. The Transceiver cell (gray) has a transgene for constitutive expression of GFP-sensing synNotch receptor with a tetracycline-controlled transactivator (tTA) intracellular domain and two transgenes with tTA-responsive elements (TRE) activating GFP ligand and mCherry (see Methods: Constructs). (C) Micrographs demonstrating *in vitro* signal propagation over time at the indicated cell densities. Bright field (grayscale) is overlaid with GFP signal (green) and nuclear infrared fluorescent marker expressed in sender cells (purple). A single propagation focus is shown, isolated from a culture well with multiple foci. Scale bar 100 *µ*m. See Supplemental Movies 1-3 for time-lapse movies. (D) Propagation radius (*r*_prop_) over time for three cell densities (*n* = 5, mean ± s.d.). (E) Cell-level mathematical model of Transceiver signaling. A sender cell (left) presents GFP ligand *s* (green ellipsoid and triangles) to a Transceiver cell (center, “*i*”). Ligand from cell i’s neighbors (“*j*”) activates SynNotch receptors (blue). Activated SynNotch stimulates production of ligand and a reporter *r* (red ellipsoid) after a time delay *τ*. Production is regulated by cell density. The ligand s also inhibits its production (“*cis*-inhibition”). Ø indicates degradation. (F) *In silico* simulation of transceiver signaling demonstrating GFP ligand (green) propagation from a sender cell (purple) at cell densities of 1x, 2x, and 4x. Scale bar 125 *µ*m. (G) Propagation velocity (*r*_prop_) *in vitro* for densities of 1x, 2x, and 4x. (H) Strip plot of propagation velocity *in vitro* (black dots; horizontal line indicates mean) and *in silico* (blue diamonds) at indicated cell densities. **p< 0.01, two-sided Mann-Whitney-Wilcoxon test.

We implemented this circuit *in vitro* in a mouse L929 fibroblast cell line and analyzed propagation dynamics while varying cell culture density. To implement the circuit, we generated Transceiver cells that express an anti-GFP receptor activating transcription of the GFP ligand. To do so, three transgenes were stably integrated (Figure **2**B): the first constitutively expresses a synNotch receptor with an anti-GFP nanobody (*α*GFP) as the extracellular domain and the transcription factor tetracycline transactivator (tTA) as the intracellular domain. The second expresses membrane bound GFP (synNotch cognate ligand) under control of the tetracycline responsive element (TRE) promoter. Finally, for tracking purposes, a transgene that expresses TRE-driven cytosolic mCherry as a readout of synNotch activity and constitutively expresses the blue fluorescent protein tagBFP to mark transceivers. We generated several clones of L929 cells where these transgenes were all integrated into the genome. Transceiver clone function was evaluated by co-culturing with sparse sender cells and performing high-magnification time-lapse imaging centered on individual sender cell foci.

The area of GFP fluorescence was calculated using a semi-automated image analysis workflow (see Methods: Image analysis). At a cell density of 1250 cells/mm^2^ (100% confluent, or “1x”), transceiver fibroblasts in contact with a sender cell express the GFP ligand, triggering a propagating wave of transceiver signaling that travels outward through the cell monolayer (images in Figure **2**C; see Supplementary Videos 1-3). The sizes of *n* = 5 signaling discs were quantified by calculating the radius of propagation *r*_prop_ (Figure **2**D), resulting in a mean velocity of d*r*_prop_/d*t* = 0.131 ± 0.009 mm/day at 1x confluent density. When initial cell density is increased, wave velocity slows to 0.085 ± 0.014 mm/day at an initial density of 2x (2500 cells/mm^2^) and 0.026 ± 0.011 mm/day at an initial density of 4x (5000 cells/mm^2^). Similar trends were observed for all clones generated (data not shown). Thus, this synthetic lateral propagation circuit can generate signaling waves whose velocity is responsive to cell density.

To gain additional insight, we devised a computational model of synNotch signaling with density-dependent attenuation, and used it to study the impact of density-dependent signal transduction on signal propagation in a multicellular sheet. To model the multicellular sheet, we use a fixed lattice of cells, a framework used extensively to study Notch-mediated patterning^49–54,70^. In our model, each cell occupies a region on a hexagonal lattice. For a given density, the area of each hexagon on the lattice is set equal to the average area of cells in a confluent monolayer *in vitro*. As depicted in Figure **2**E, each cell contains a system of chemical reactions that model synNotch signal transduction (see Methods: Mathematical modeling, Equations (3)). A given cell (*i*) in direct contact with its neighbors (*j*) can express both the signaling ligand (*s*) and the reporter protein (*r*). Cells either express the ligand constitutively if they are senders or upon synNotch stimulation if they are transceivers. To model cytoplasmic projections, which are known to affect Notch-mediated patterning systems^49,71,72^, contact strength is weighted by cell-cell distance on the lattice (weights visualized in Figure S5A).

Following ligand-induced receptor activation, the recipient cell responds after a time delay (*τ*) corresponding to transcription, translation, and membrane trafficking of the ligand. Finally, ligand expression in *cis* decreases a transceiver cell’s capability to sense ligand expressed by neighbors in *trans*, a phenomenon termed *cis*-inhibition that is observed to regulate endogenous and synthetic Notch signaling^51^. Please see Figure S5B for an example of sender, receiver, and transceiver activation dynamics *in silico*. Changes in cell density are modeled by changing the size of all cells equally. At a higher density, for example, cell size is reduced to occupy less area while preserving the hexagonal lattice (Figure S5C, inset images). The density-dependence of signaling is phenomenologically modeled by multiplying the amount of ligand involved in signaling by a coefficient that encodes the efficiency of signaling. This coefficient decays as cell density exceeds confluence (Figure S5C, blue curve) and was parameterized by comparison with the propagation data in Figure **2**.

With this *in silico* model of density-dependent, signal-propagation circuit, we sought to simulate propagation on a monolayer lattice at different densities. To do so, we initialized a 50 × 50 hexagonal lattice of transceiver cells with a single sender cell. At time *t* = 0, the sender cell begins expressing the signal and Equations (3) are integrated forward in time (see Methods: Mathematical modeling and Supplementary Text for details of mathematical modeling). We perform the simulation at cell densities of 1X, 2X, and 4X confluence (1250, 2500, and 5000 cells/mm^2^). In the simulations, a wave of activation begins propagating outwards from the Sender at a speed that depends on density (selected time-points rendered in Figure **2**F; see Supplementary Video 4). The area of propagation was quantified as the amount of lattice area occupied by Transceivers expressing ligand at a level greater than the promoter threshold *k* (see 1 for values of all parameters used in simulation). Wave velocity was calculated similarly to the *in vitro* case. As shown in Figure **2**G, in the 1x condition propagation area begins to increase after a time delay *τ* = 0.3 for signal production and then continually rises. By itself, the time-scale of simulation is dimensionless and thus has no intrinsic units. Thus, we show time in units of *τ* as a characteristic time-scale. At 1x density, *in silico* wave velocity (calculated similarly to *in vitro*) is 0.103 mm/*τ* and drops to 0.055 mm/*τ* and 0.009 mm/*τ* at 2x and 4x densities, demonstrating that the attenuation of signaling at higher cell densities is sufficient to explain the slower speed of Transceiver propagation. Figure **2**H compares the mean velocities of the experimental system and computational model. In both cases, increasing cell density above 100% confluence has a substantial suppressive effect on the velocity of the wavefront.

Collectively, these results show that the signaling dynamics of a synthetic circuit can be modulated via cell density. They further suggest that cell density could be used as a mechano-chemical control mechanism to dynamically pattern synthetic tissues without requiring the engineering of additional biochemical circuit components.

### Signal propagation reaches self-limiting regimes due to cell population growth

Cells in culture grow and divide, thereby increasing cell density over time. How does population growth impact signaling behavior over longer time courses? Figure **3**A shows an example of a seven-day time course in which transceivers were co-cultured with senders at a plating density of 1250 cells/mm^2^ (1x). As shown in the GFP channel images, transceivers begin propagating signal by Day 1 of growth, and the signaling wave propagates outwards over the first three days. By Day 4-5, however, propagation speed and overall signal intensity start to decline, and by Day 7 GFP expression is almost fully suppressed. This resulted in self-limiting propagation with a characteristic diameter of 0.5 mm. Interestingly, transceivers that express and then down-regulate GFP signal continue to express the mCherry reporter at Day 7 (mCherry channel), suggesting slower degradation kinetics. Importantly, cells re-plated at 1x density after a seven-day time course are still capable of propagation after re-plating, suggesting that GFP down regulation is not a result of decreased cell health or death during long-term culture (Figure S6).

**Figure 3:**
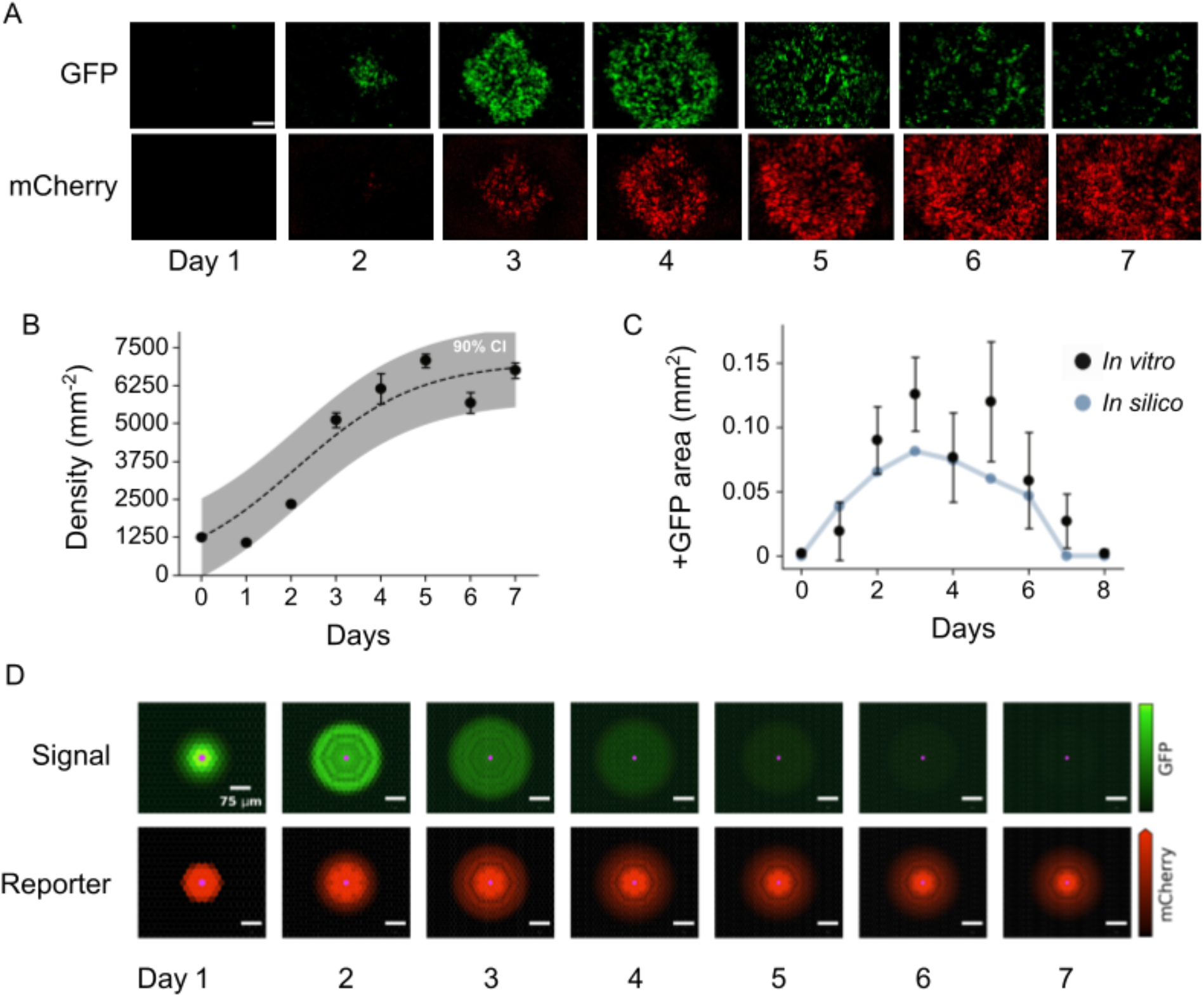
Cell population growth over time leads to self-limiting activation. (A) Propagation and attenuation of signal over a seven-day time-course. Fluorescence micrographs were taken of an isolated propagation focus from a sender:transceiver co-culture plated at a density of 1250 cells/mm^2^ (1x). GFP produced by senders and activated transceivers is shown in green, and mCherry (reporter for synNotch activation in transceivers) is shown in red. Signal begins propagating in a disc until around day 4. From then, GFP levels decrease and are mostly decayed by day 7. The reporter remains as a record of past activation, possibly due to slower degradation kinetics. Scale bar 100 *µ*m. (B) Graph of cell density over time. Black dots are the mean and standard deviation (*n* = 3) of cell density from sender-transceiver co-cultures, measured by automated cell counting. The dashed line shows the logistic growth equation with estimated parameters (90% confidence interval in gray). (C) Quantification of the propagation radius (*r*_prop_) over time. *In vitro*, 1:100 co-culture of senders:transceivers was plated at 1x density, and the areas of *n* = 5 individual foci were imaged daily. *In silico*, an 80 × 80 lattice of transceivers with one sender was simulated from an initial density of 1x. Activated area was measured as in Figure 2 for *in vitro* (black dots, mean ± s.d.) and *in silico* (connected blue dots). Cell density is increased over time by reducing cell size to account for population growth as estimated in (B). Simulation time-scale was parameterized using the estimated generation time (see Results and Methods: Mathematical modeling). Population growth thus appears sufficient to explain the experimentally observed dynamics of ligand activation followed by attenuation. (D) Rendering of model simulation. The model consists of a hexagonal lattice with one sender cell (purple) surrounded by transceivers at 1x density. GFP ligand (first row, green) and mCherry reporter (second row, red) concentrations in transceivers are rendered at daily time-points.

Given the negative correlation between cell density and signaling, we hypothesized that the decrease in GFP signaling at later times is due to an increase in cell density secondary to cell proliferation. To test this hypothesis, we counted cell numbers over the time course in culture. Transceivers were plated at initial densities of 1250, 2500, and 5000 cells/mm^2^ (1x, 2x, and 4x), and n = 3 cell counts were performed daily for seven days using an automated cell counter. Results for the 1x condition shown in Figure **3**B demonstrate how cell density (black, mean ± s.d.) increases over time in a sigmoid fashion and begins to plateau by 4-5 days of culture. By day 3 - around the same time signaling begins to shut off in Figure **3**A - the culture achieves a density of ≥ 3750 cells/mm^2^ (3x) which was found to be inhibitory to transceiver signaling (Figure **1**E, 3x and 4x density). These results supported the hypothesis that in long-term culture, cell proliferation can shut off signaling in previously activated transceivers if the culture achieves cell densities that are not conducive to cell-cell signaling.

We then used our computational model to determine if density-induced repression is sufficient to replicate signaling attenuation in long-term cultures. We model population growth as a logistic growth process. The logistic growth equation is a common sigmoidal growth model in which a species will proliferate exponentially until it approaches the maximum population density its environment can support (termed the “carrying capacity” *ρ*_max_), at which point density asymptotically approaches *ρ*_max_. Like exponential growth, logistic growth starts at an initial population density *ρ*_0_ and has an intrinsic growth rate *g*. We applied a maximum-likelihood estimation (MLE) procedure to our cell count data and estimate that *g* = 0.616 days^−1^ (90% CI: 0.539 - 0.711 days^−1^) and *ρ*_max_ = 7337 cells/mm^2^ (90% CI: 6954 - 7771 cells/mm^2^). In Figure 3B, we plot the fitted logistic growth equation (black dashed curve, 90% CI in gray), which shows good correspondence with the observed growth data. Please refer to Methods: Statistical analysis and Figure S7 for detailed MLE procedure and results. Given this parameterization, the generation time of cells is ln 2/g = 1.125 days (27.0 hrs). Under the simplifying assumption of a constant protein removal rate, we can equate the generation time to the ligand half-life, allowing us to convert between simulation time and real units (see Mathematical modeling for a detailed discussion).

Transceiver signal propagation was then simulated with logistic growth of the population density over time. We generated an 80 × 80 lattice of transceivers with one sender at the center at an initial density of 1x and ran the simulation for 8 days. Changing population density in our system is modeled by reducing the area of cells on the lattice rather than by explicit cell division, mitigating computational complexity (see Methods: Mathematical modeling for details). The propagation radius was then measured as in Figure **2**G and plotted over time in Figure **3**C (blue) alongside the radii of *n* = 5 *in vitro* propagation foci (black, mean ± s.d.) under the same experimental conditions as Figure **3**A. The simulated dynamics of signaling are similar to that of the *in vitro* results. In particular, the *in silico* model also exhibits an early period of 3 days of activation characterized by GFP and mCherry production and expansion of the propagation disc, followed by a period of signal attenuation during which GFP expression falls and eventually reaches levels similar to day 0. Renderings of the simulation at daily time-points are shown in Figure **3**D. To model the lingering nature of the mCherry reporter, mCherry was modeled under the same activation kinetics as GFP but with 10 times slower degradation kinetics (see Methods: Mathematical modeling).

These results demonstrate that cell density changes are sufficient to explain the observed attenuation of transceiver activation and signal propagation. Overall, we show using experimentation and mathematical modeling that signal propagation through a proliferating transceiver population can have a transient, self-limiting nature consistent with a density-induced attenuation of synNotch activity.

### *In silico* exploration of growth parameters reveals distinct phases of activation explained by a critical density

Using the computational model, we then investigated the generative possibilities of this mechanochemically coupled circuit. So far, in the system, we have observed that the speed of propagation can be controlled via initial cell density (Fig. 2), and also over time due to cell proliferation (Fig. 3). This observation lead to identify initial cell density, and cell proliferation as two variables affecting patterning outcomes. In order to define the space of achievable qualitative and quantitative phenotypes for the density-modulated Transceiver signaling circuit, we simulated circuit behavior for different values of the initial density *ρ*_0_ and proliferation rate *g* and generated a phase diagram of signaling phenotypes. For each parameter combination, a 50 ρ 50 lattice of transceiver cells and one Sender were simulated for 9.5 days. We observe signaling behavior that falls into three categories. Either signal begins to propagate or not, and if propagation begins, Transceivers either all become deactivated by the end of the simulation or remain active to some extent. Parameter sets thus fall into three categories, or “phases”: attenuated, self-limited, or unlimited propagation. For classification, signaling dynamics at early and late time points are used. Parameter sets are labeled “attenuated” if the initial signal production rate 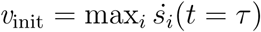 is below a chosen threshold *v*_thresh_ = 0.25 (Figure S8B). If the initial production rate is above this value, they were labeled s”self-limited” if all Transceivers become inactivated by the end of the time-course and “unlimited” otherwise. As above, a transceiver *i* was considered activated based on the amount of expressed ligand (*s*_*i*_ > *k*; see Table 1).

**Table 1:** Modeling parameters. ^†^Expressed in dimensionless time-units; corresponds to 0.43 *t*_*d*_, where *t*_*d*_ is the doubling time of the cell line.

| Parameter | Meaning | Value used |
| --- | --- | --- |
| $\alpha$ | Ligand production rate when induced | 3.0 |
| $k$ | Signaling threshold | 0.02 |
| $p$ | Cooperativity (ultrasensitivity) of signaling | 2.0 |
| $\varepsilon$ | Strength ligand-receptor inhibition in *cis* | 1.0 |
| $\tau$ | Time delay for production | 0.30<sup>†</sup> |
| $\gamma$ | Reaction rate constant of $R$ relative to $S$ | 0.1 |
| $m$ | Sensitivity of synNotch signaling to density | 1.0 |
| $v_{\text{thresh}}$ | Threshold for initial ligand production | 0.5 |

Each of the three behavioral phases corresponds to a discrete region of the phase diagram. In the dark blue region of Figure **4**A, Transceiver activation persists throughout the time-course and propagation is unlimited. In the blue region of limited propagation, the wave of activation initiates but becomes fully attenuated at some point during the time-course. In this regime, Transceiver activation is limited in time and space, and the maximum area achieved during the time-course depends on both *ρ*_0_ and *g* (Figure **4**B). Finally, in the gray region, no activation occurs and signaling is fully attenuated above a critical density (*ρ* > *ρ*_crit_ ≈ 3.25) that does not depend on *ρ*_max_ (Figure S5). Therefore, these phases represent different collective signaling dynamics (Figure **4**C). We noticed that the observed phases correspond to the growth curve lying below *ρ*_crit_, crossing *ρ*_crit_, or lying above *ρ*_crit_ (Figure **4**E); this is due to the monotonic nature of logistic growth and the effect of each parameter in the logistic equation (Figure **4**D). In particular, the attenuated propagation phenotype can be accessed above a specific initial cell density, regardless of proliferation; for the other 2 phenotypes where propagation is observed, whether the systems displays a self-limiting behavior or propagates in an unlimited fashion can be controlled via controlling proliferation even for cells plated at the same initial density. In sum, we identified three dynamical behaviors in silico that can emerge in cells harboring the same genetic circuit, and these behaviors can be accessed by manipulating parameters of cell proliferation such as intrinsic proliferation rate g and initial cell density *ρ*_0_.

**Figure 4:**
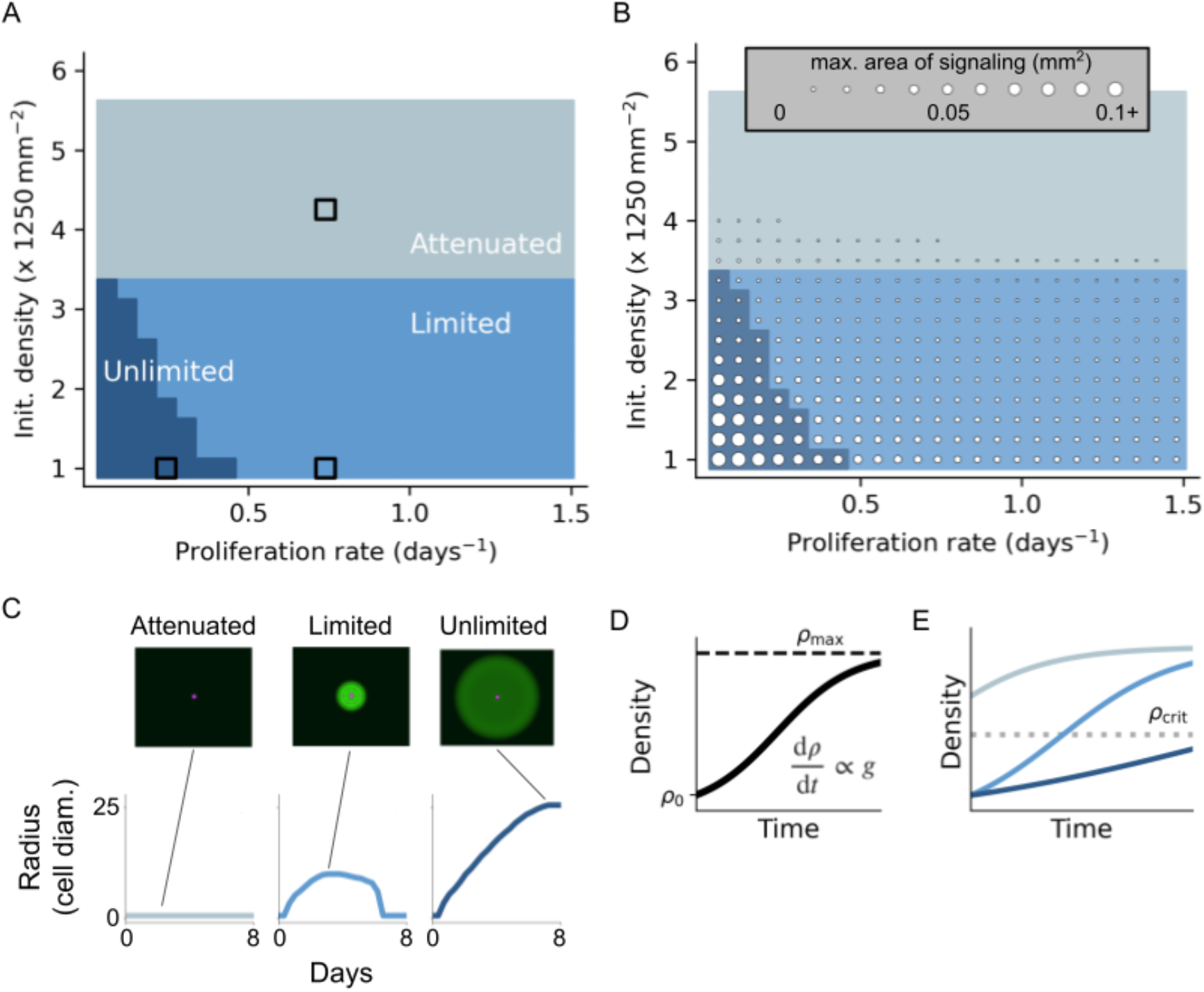
Population growth determines the signaling behavior of transceivers. (A) Phase diagram showing the transceiver propagation phase as a function of initial cell density and proliferation rate. For each simulation, a 50 × 50 lattice of Transceiver cells and one Sender were simulated with different values of the growth parameters *g* (intrinsic proliferation rate) and *ρ*_0_ (initial density) for 9.5 days. Parameter sets were then classified into distinct phases based on signal propagation behavior at early and late time-points: unlimited (dark blue), self-limited (light blue), or attenuated (gray) propagation. In the unlimited phase, the ligand production rate is too low at early time-points to trigger activation. In the self-limited phase, activation occurs but all activated Transceivers become deactivated by the end of the simulation, and in the unlimited phase, some activated cells remain. Black squares are example parameters for which dynamics are shown in (C). (B) Phase diagram annotated with circles showing the maximum area of the propagation disc over the whole time-course. The disc reaches negligible sizes in the attenuated phase and persists at the end of simulation time in the unlimited phase. However, in the self-limited phase, propagation reaches a finite area. Due to the reporter’s slow degradation kinetics, transceivers “remember” prior activation. Thus, the maximum area of propagation could be tuned by modifying the parameters of growth and subsequently maintained in molecular memory. (C) Example time-courses and simulation renderings for each phase. Parameters correspond to outlined squares in (A). The radius of the activation radius is shown over time, and above each graph is shown a simulation rendering at the time of maximum disc area. Time-course was sub-sampled for clarity. (D) An example of logistic growth. Population density begins at an initial density *ρ*_0_ at *t* = 0 and increases sigmoidally at an intrinsic rate *g*, approaching the asymptotic carrying capacity *ρ*_max_. (E) Cell density over time for the three behavioral phases. The three phases can be understood in relation to a threshold density *ρ*_OFF_ (dotted line) above which signaling becomes attenuated. Attenuated tissues are more dense than *ρ*_OFF_, unlimited tissues are less dense than *ρ*_OFF_, and self-limited tissues cross the threshold during the time-course.

### Growth rate-modulating drugs push Transceivers into different phase regimes

According to the computational model, decreasing the intrinsic rate of proliferation (*g*) should greatly extend the amount of time spent at densities permissive to signaling and therefore shift our system from a regime of self-limited propagation into a regime where propagation is virtually unlimited (from the light blue region to the dark blue region in the Figure **4**A phase diagram). Conversely, increasing the proliferation rate within the self-limited region should decrease the radii of propagation foci (light blue region in Figure **4**B). Finally, increasing the initial plating density (*ρ*_0_) of the culture greater than 1250 cells/mm^2^ (1x) should reduce the size of propagation foci.

To test these predictions *in vitro*, we performed propagation experiments in conditions that perturb cell proliferation. Cells were cultured in the presence of Y-27632^73^, a small molecule ROCK inhibitor (RI) that limits proliferation^74–76^, or fibroblast growth factor 2 (FGF2), a growth factor that stimulates proliferation^77–79^. Sender:transceiver co-cultures were plated at 1x, 2x, and 4x densities, cell density was counted daily (*n* = 3) for seven days, and the effect on the intrinsic proliferation rate *g* was estimated similarly to Figure **3**. Figure **5**A shows growth curve data from the 1x plating density, color-coded by drug treatment (colored circles, mean ± s.d.; see Figure S9 for 2x and 4x initial densities). Treatment with 250 ng/mL FGF2 (purple) accelerated proliferation by a factor of 1.49 to 1.09 days^−1^ (90% CI: 0.847 to 1.45 days^−1^) compared to the untreated condition (yellow), while 50 *µ*M RI (green) slowed proliferation by a factor of 0.19 to 0.141 days^−1^ (90% CI: 0.111 to 0.169 days^−1^). The resulting best-fit growth curves are shown as solid curves in Figure **5**A. These measured proliferation rates are shown in Figure **5**B (circles, mean ± 90% CI) superimposed on the phase diagram of behavior predicted by the computational model. In this model, RI treatment (green) should push the system into the dark blue region of “unlimited” propagation that does not shut down by day 8, while FGF2 treatment (purple) should remain in the light blue region of “self-limited” propagation (respectively, left and right in the X-direction in the phase diagram). Of note, neither drug was found to modulate synNotch activity *per se* (see Figure **1**D and Figure S8G). See Figure S9 for full results of MLE.

**Figure 5:**
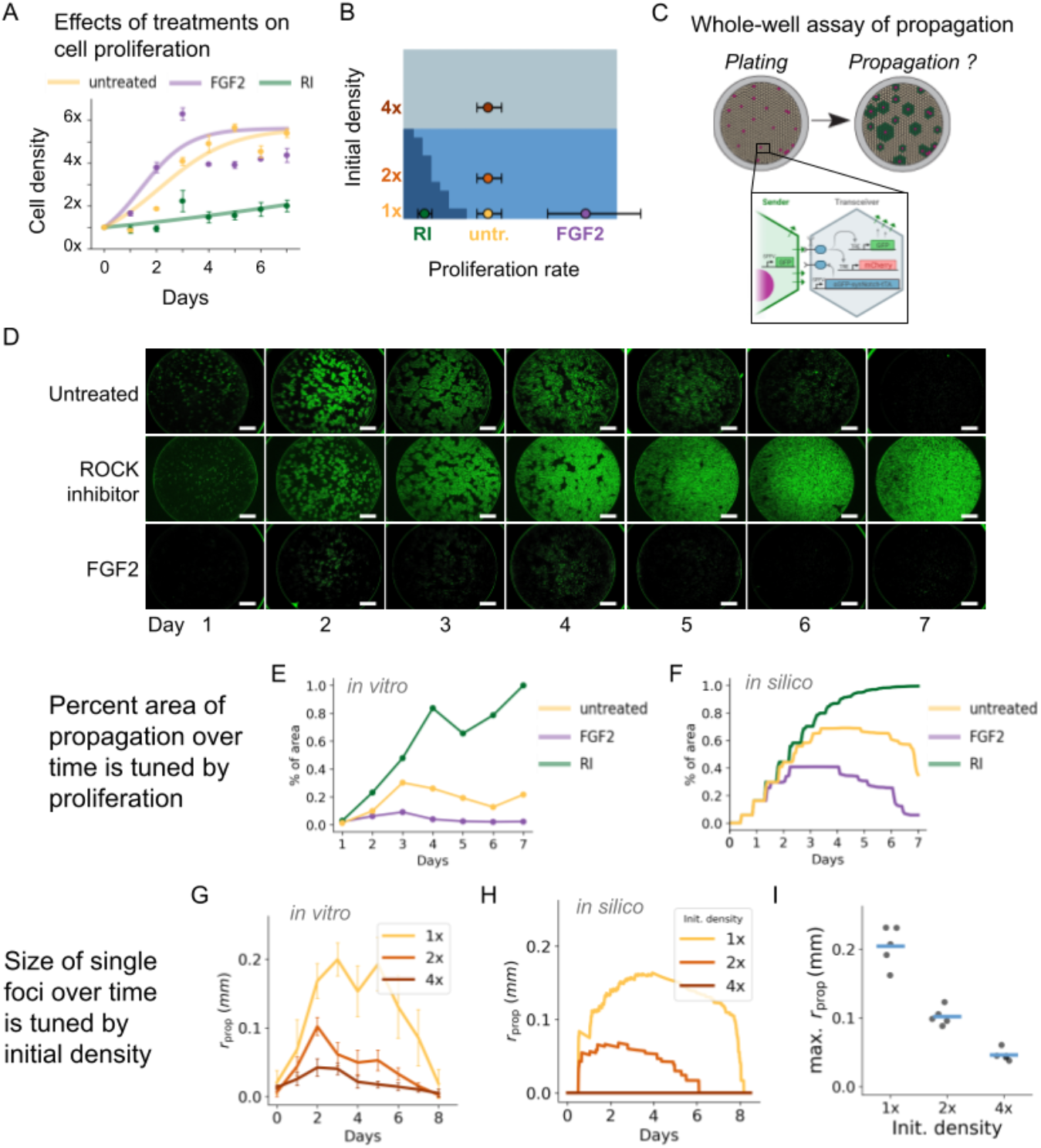
*In vitro* control of Transceiver activation area by manipulating either population growth rate or initial population density. (A) Time-series of cell density under growth-modulating drug conditions: Untreated (yellow), FGF2 (violet), or ROCK-inhibitor (RI; green). Cell count data (circles, mean ± s.d.) were used to parameterize the logistic growth equation (solid lines). FGF2 induces faster population growth, while RI dramatically slows growth. “Untreated” sample reproduced from Fig. **3**B. (B) Experimental perturbations alter the predicted behavior of transceivers. Phase diagram of transceiver propagation behavior as a function of proliferation rate (*g*) and initial cell density (*ρ*_0_) (see Figure **4**A). Circles indicate *g* (estimated mean and 90%CI) and *ρ*_0_ for real *in vitro* perturbations. An increase in *ρ*_0_ from 1x (yellow) should cause smaller foci (2x, orange) and eventually full attenuation of signaling (4x, red). Alternatively, slowing down growth should cause fast, uncontrolled propagation (low *g*, green) while speeding up growth should cause faster attenuation (high *g*, purple). (C) Schematic of the whole-well propagation assay. A co-culture of senders (purple) and transceivers (brown) is plated in a culture well at time *t* = 0. Each sender acts as a propagation focus (inset diagram), and the amount of ligand produced in the well (green) over time is measured by fluorescence imaging. This assay allows quantification of propagation without isolation of single propagation foci. (D) Time-series micrographs of 1:100 sender:transceiver co-cultures plated at 1x density (1250 cells/mm^2^) under various drug treatments. Senders and activated transceivers produce GFP (green). Compared to untreated, RI-treated wells propagate strongly throughout the time-course, while FGF2-treated wells propagate much weaker (days 1-4) and shut off earlier (day 5-6 instead of 7). Scale bar 1 mm. Note the well border produces a circular artifact. (E) *In vitro* and (F) simulated results for the assay in (D), shown as the percentage of the well covered by GFP fluorescence over time. (G) *In vitro* and (H) simulated propagation radii (*r*_prop_) of single foci over time for different initial densities. Error bars in (G) indicate mean ± s.d. (I) Peak propagation distance *in vitro* for different initial densities (*n* = 5, blue indicates mean).

To test these predictions, we devised a “whole well” assay of signal propagation. Figure **5**C shows a schematic of this assay in which senders (purple) and transceivers (brown) are initially plated in a 1:100 ratio (top left diagram). Each sender (inset diagram, bottom) locally triggers a wave of GFP signaling (green), and in permissive conditions, these foci expand and fuse over time to occupy a significant percentage of the surface area of the culture well (top right diagram). This assay allows us to quantify transceiver propagation in cases where isolation of individual propagation foci is challenging, such as when propagation is so efficient that adjacent foci in the well rapidly fuse.

Using this assay, we tracked GFP fluorescence in whole culture wells over a seven-day timecourse under different drug treatment conditions. Figure **5**D shows that in the absence of drug treatment, transceivers initially produce GFP ligand and subsequently shut off expression by day 6-7 (first row, “untreated”), as in the single-foci experiments. In the presence of RI, however, GFP ligand propagation proceeds virtually indefinitely and fills the entire culture well (second row, “ROCK inhibitor”). Conversely, FGF2 treatment causes a more subtle activation of GFP ligand, followed rapidly by GFP depletion (third row, “FGF2”). The percentage of the well covered by GFP fluorescence is shown in Figure **5**E (untreated: yellow, RI: green, FGF2: purple). We populated the morphospace also with the other combinations of proliferation rate/initial density combinations, the results follow similar trends and are shown in Fig. S10C,D.

Figure **5**F shows computational simulations of these scenarios. For each experimental condition, a 150 × 150 hexagonal lattice was randomly seeded with senders and transceivers in a 1:100 ratio (*n* = 10 replicates) and numerically integrated over seven days using the population growth parameters fitted via MLE and a shorter cell-cell contact distance (see Methods: Mathematical modeling). In the absence of treatment (yellow), a majority of the lattice quickly activates, followed by a gradual decline. In the presence of RI (green), propagation instead persists throughout the time-course, eventually saturating the lattice. In contrast, FGF2 treatment (purple) speeds up the progression from propagation to attenuation, causing lower levels of activation. Thus, our model corroborates the experimental evidence and demonstrates that the effects of RI and FGF2 treatment on cell proliferation (inhibition and activation, respectively) are sufficient to explain the starkly divergent outcomes of signaling activation between these conditions. We also note that the experimental propagation area was found to be smaller than predicted by modeling (contrast the peak activation in the Untreated and FGF2 conditions in Figure **5**E and Figure **5**F), even with a shorter contact distance. Such a deviation could be a result of the lower signal-to-noise ratio when imaging at low magnification, where low levels of fluorescence are more difficult to discern and isolate from background. Additionally, drug treatments appear to induce changes in signaling behavior within the first two days of growth, earlier than predicted by modeling (contrast the separation between curves in Figure **5**E and Figure **5**F). In addition to altering population growth, is therefore these drugs may affect other factors, such as the production rate of ligand and/or the process of fusion between adjacent signaling foci.

Alternatively, the Y axis of the phase diagram can be traversed by changing the initial cell density. Single-foci propagation assays were performed as described above, *in vitro* (*n* = 5) and *in silico*, starting from initial densities of 1x, 2x, or 4x confluent cell density. Figures **5**G-I show that the behavior of the propagation follows the predictions of the model. In Figure **5**G (*in vitro*; mean ± s.d.) and Figure **5**H (*in silico*), the propagation disc is smaller and deactivates faster at an initial density of 2x. At an initial density of 4x, there is no activation *in silico* and very little activation above baseline *in vitro*. Thus, as initial density increases, the critical attenuation density is reached more quickly and there is less time for signal to spread. Due to the persistence of mCherry after transceiver deactivation (see Figure **3**A), an important property of transceiver patterning is the maximum size of the activation disc before deactivation. As shown in Figure **5**I, the initial density strongly determines the maximum area of the propagation disc (*n* = 5 replicates in black; blue lines indicate means). These perturbation experiments support the idea that population density dynamics are predictive of transceiver patterning. They also demonstrate the feasibility of treating two parameters of growth - initial cell density and the intrinsic rate of cell proliferation - as control parameters of patterning.

### Tissue-scale cell density gradients generate spatial signaling activation gradients and kinematic waves

In order to define distinct regions of differentiation, embryonic tissues regulate the spatial distribution of chemical morphogens^46,80,81^. Recently, chemical gradients have been engineered to direct spatial differentiation using exogenous morphogens^32,82^. Here, we instead set out to engineer spatial information in a non-genetic fashion in the form of cell density gradients, which can be decoded by transceivers into distinct spatial domains.

We hypothesized that a spatial gradient of cell density can elicit a gradient of GFP ligand. In Figure **6**A, a phase diagram at an early time-point (left, computed as above at *t* = 2.7 days) shows that the initial cell density primarily determines whether a region of tissue is signaling or not signaling. Thus, we reason that a culture well inoculated with senders and transceivers in a gradient of cell density (top right) should become activated in areas where density is within the optimal range (*ρ*_ON_ < *ρ* < *ρ*_OFF_) and stay inactive in areas where density is above the optimal range (*ρ* > *ρ*_OFF_). Figure **6**A, bottom right, shows a computational example where an initial density gradient causes a gradient in the predicted amount of GFP ([GFP]_SS_, shown in green).

**Figure 6:**
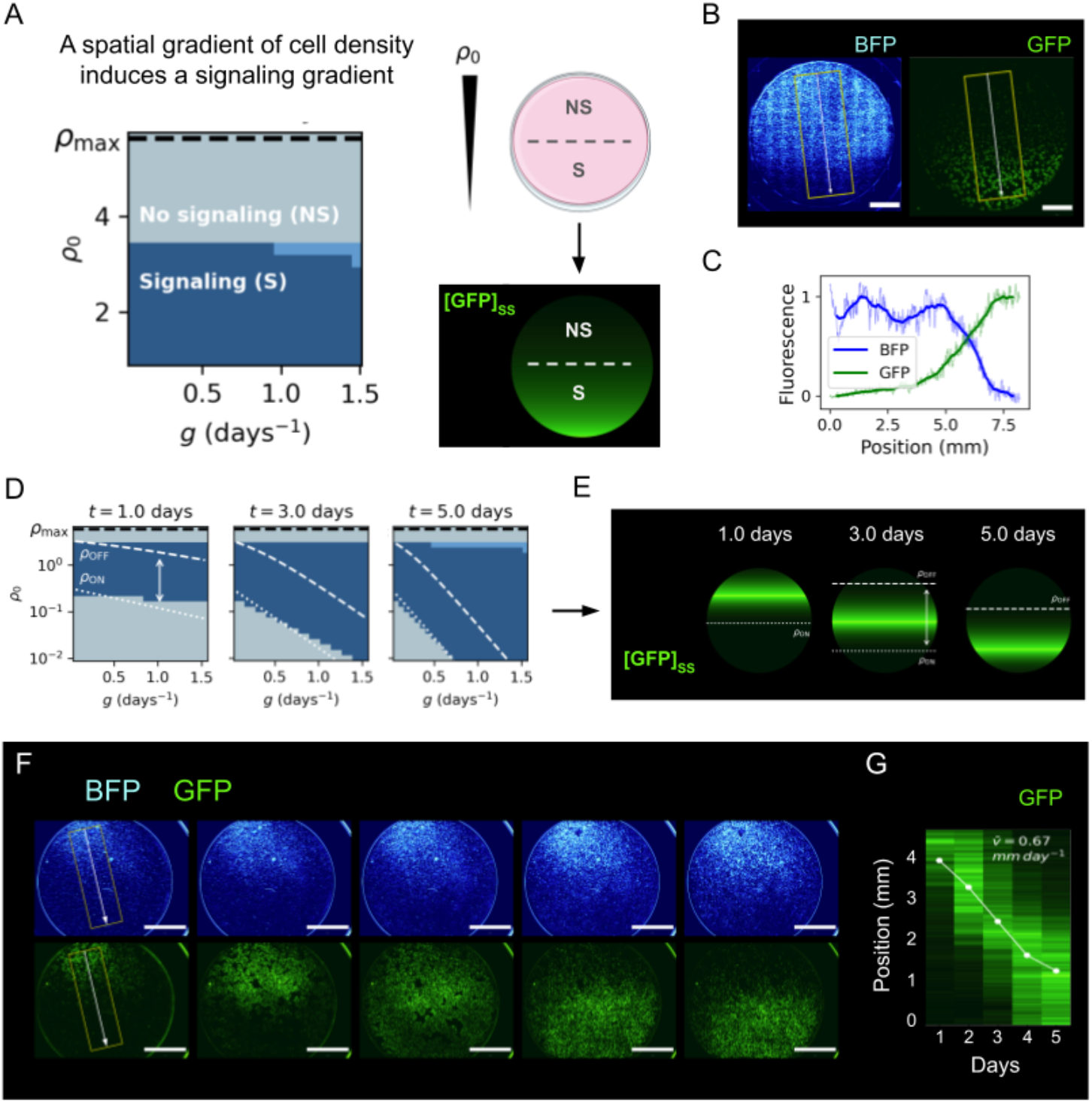
Spatial gradients of cell density produce long-range activation gradients and kinematic waves. (A) Modeling suggests that at early time-points, transceiver activation is determined primarily by the initial density *ρ*_0_ (left, *t* = 2.7 days). Spatial variation in signaling, therefore, could be achieved by biasing the initial seeding of the cell culture well to create a spatial gradient of density (top right). If the the top half of the well is too dense for signaling while the bottom half is optimal, distinct regions of GFP signaling/no signaling should develop (bottom right). Green represents *[GFP]* at steady-state ([GFP]_SS_). (B-C) *In vitro* density and signaling gradients. (B) Stitched epifluorescence micrographs of a culture well inoculated with a sender:transceiver co-culture (1:100 ratio) in a spatial gradient of initial density (average of 2x) and imaged at 64 hrs (2.7 days) of culture. Scale bar 2 mm. TagBFP is constitutively expressed by Transceivers and acts as a cell density readout. Boxed region quantified in (C). (C) Fluorescence profiles showing the anti-correlated BFP and GFP gradients. (D-E) *In silico* modeling predicts long-range kinematic waves over time. (D) Phase diagrams at three time-points with logarithmic y-axes. The density range between *ρ*_ON_ and *ρ*_OFF_ (white dotted and dashed lines) is optimal for signaling. (E) Given an initial density gradient, different areas of the well could enter/exit the optimal range at different times, creating a virtual (kinematic) wave of activation. Green is [GFP]_SS_. (F-G) A synthetic kinematic wave generated by a density gradient *in vitro*. (F) Epifluorescence micrographs of a culture well seeded with 1:100 senders:transceivers in a spatial gradient of density (average of 1x) and imaged daily. From days 1-4, a wave spreads downwards as different regions become activated. Scale bar 2 mm. The boxed region is quantified in (G). (G) Spatial profile of GFP fluorescence in (F) over time. White dots show the mean wavefront position, which has a mean velocity of 0.67 mm/day.

To test this prediction, we established an *in vitro* system where a a 1:100 mixture of senders and transceivers was seeded in a tissue culture well. Based on the number of cells seeded, the density was 2x confluence (2500 cells/mm^2^) on average, but the initial seeding was biased towards one end of the well, forming a gradient of cell density. We then imaged the constitutive blue reporter (tagBFP) and the GFP ligand as a proxy of cell density and signaling activation, respectively. As shown in Figure **6**B, at 2.7 days of culture the GFP output of the transceivers is patterned along the well in a way that recapitulate the cell density gradient. The yellow boxes indicate the region used to quantify the BFP and GFP gradients shown in **6**C. Thus, spatial patterns of gene expression can be established with this circuit given the cell-density dependency, via establishing cell density gradients.

As with uniform cell density, we reason that gradients of cell density can produce rich signaling behavior over time due to the dynamics of cell population growth. In particular, because signaling activation occurs in our system within an optimal range of density (see **1**E), we hypothesized that at lower densities, a spatial gradient of cell density could cause a virtual wave of activation across the well. Due to population growth, a region of the culture well that begins the time-course too sparse for activation (*ρ* < *ρ*_ON_) over time will enter the optimal range (*ρ*_ON_ < *ρ* < *ρ*_OFF_) and eventually exit this range (*ρ* > *ρ*_OFF_). In Figure **6**D phase diagrams at three consecutive timepoints of simulation are annotated with the critical densities *ρ*_ON_ and *ρ*_OFF_, plotted respectively as white dotted and dashed lines. Over time, regions of the well with different initial densities (shown on a logarithmic Y-axis) should enter the optimal range between the two lines at different times. As shown using an example gradient in Figure **6**E, this staggered activation creates the appearance of a large wave spreading through the well. However, this wave is in fact only virtual, or “kinematic,” meaning each region is turning on and off independently based on its local cell density.

Our results in Figures **6**F and **6**G demonstrate a kinematic wave generated by an *in vitro* density gradient of sender and transceiver cells (1:100 ratio). The upper region of the well in Figure **6**F starts at an efficient density for signaling but eventually becomes too dense. In contrast, the bottom region of the well is initially too sparse for signaling but enters the optimal range of density after 3-4 days in culture. Although these regions are many millimeters apart (scale bar 2mm), their entry and exit into the optimal range are staggered, creating the appearance of a wave moving downwards. The data in the yellow box were used to calculate the mean wavefront position as shown in Figure **6**G, and the mean wave velocity was found to be 0.67 mm/day. This speed is 5.1 times faster than the speed of direct cell-to-cell propagation measured at 1x confluent density (see Figure **2**H) and roughly twice the speed of directed fibroblast motility^83^, making signal transduction and bulk transport unlikely causes of this phenomenon. Collectively, these results show that we can create spatial and spatio-temporal gradients of signal activation by generating gradients of cell density and exploiting the dynamics of population growth.

## Discussion

The astonishing diversity of tissue patterning and morphology in our own bodies underscores the critical importance of genetic circuits that operate in an ever-changing mechanical environments, and provokes the fundamental question of whether and how mechano-chemical circuits expand achievable behaviors, compared to chemical only circuits, to achieve some of the organization that we see in mammalian multicellular tissue shapes and patterns.

Here, we discovered a new physical and non-genetic input, on synNotch, which was previously unknown (Fig. 1), cell density, and use it to build a signal-propagation patterning circuit. This circuit displays many of the hallmarks of natural patterning systems that operate in a morphologically non-inert contexts. The coupling between density and signal propagation means, in practice, that the physical state of a growing multicellular structure switches the circuit between propagating and non-propagating phases. This allows for control of behaviors such as self-limiting propagation (Fig. 2-3), propagation speed, and area dependent on cell-division parameters (Fig. 4-5), spatial gradient of activation from genetically identical cells (Fig. 6), all with this only one simple circuit. Through the coupling, the circuit locally senses the physical state (density) of a multicellular structure and the mechanical information is converted into a signaling currency, and this allows the generation of more complex behaviors.Similar control principles (signaling controlled by density) may play related roles in natural systems, allowing the control of minimal genetic patterning circuits, once evolved, via non-genetic mechanisms, to generate a variety of tissue patterning, in a manner analogous to what is shown by the density-controlled patterning observed here.

A remarkable feature of this circuit is its close agreement with predictions from a computational systems model (Fig. 2-6). Despite a lack of precise quantitative parameter values for many molecular interactions, the qualitative behaviors possible with this density-dependent multicellular circuit can be enumerated and explained from simple properties of the components and their interactions. More precise measurements could help to explain, eliminate, or exploit asymmetries and provide a better understanding of the timescales of state transitions.

Future work can explore the impact of physical variables on different synthetic gene circuits and also explore multicellular designs that explicitly exploit physical-chemical coupling to control synthetic morphogenesis using synthetic gene circuit dynamics. The physical environment can impact gene circuits both through direct mechanical input into circuit dynamics and also through altering the shape of the manifold on which reaction-diffusion circuit dynamics proceed.

The development of more complex circuit architectures that combine additional patterning capabilities (e.g. self-organizing Turing patterns, checkerboard) with dependency on additional environmental inputs (e.g. thermo-dependency, osmotic-dependency, mechanotransduction) and an ability to function on shapes that define a range of 2D and 3D tissue manifolds (in 2D in curved or confined surfaces, in 3D elongating, segmenting, branching) could enable the control of ever-increasing complexity of patterning and morphogenesis in complex growth environments. Physical-chemical coupling phenomena provide a route towards constructing synthetic circuits that can modulate progression through morphogenesis in a stepwise fashion, for example, executing new gene expression programs sequentially following the completion of a morphogenetic program, towards the construction and control of circuits of complexity similar to the ones observed in vivo.

## Methods

### Constructs

#### Constructs Design

The pHR_SFFV_LaG17_synNotch_TetRVP64 (*#*79128). The pHR_SFFV_GFPligand (*#*79129) were provided by AddGene, whereas the pHR_TRE3G_mCherry_PGK_BFP was obtained as described in the original syn-Notch paper (Morsut et al., 2016). The rest of the constructs were cloned via In-Fusion cloning (Clontech *#*ST0345). Specifically, the plasmids used for engineering fibroblasts were cloned in the pHR plasmid for lentivirus production.

#### Lentivirus Production

Lentivirus was produced by co-transfecting the transfer plasmids (pHR) and vectors encoding packaging proteins (pMD2.G and pCMV-dR8.91). Plasmids were transfected by lipofectamine LTX transfection reagent (ThermoFisher Scientific) in HEK293-T cells plated the day before in 6-well plates at approximately 70% confluence (800000 cells/well). Supernatant containing viral particles was collected 2 days after transfection and filtered to eliminate death cells and cellular debris (cut-off 0.45 *µ*m).

#### Cell Lines

L929 mouse fibroblast cells (ATCC*#* CCL-1) and HEK293 cells were cultured in DMEM (Invitrogen) containing 10% fetal bovine serum (Laguna Scientific) and tetracycline (100 ng/ml) when requested. The L929 “senders” were transduced to stably express surface GFP (GFP fused to the PDGFR transmembrane domain) and a nuclear infrared fluorescent marker (H2B-miRFP703). L929 “transceivers” were transduced with three different virions. The first virion constitutively expresses an anti-GFP antibody (Lag17) fused to a syn-Notch receptor with the transcription factor tetracycline Trans-Activator (tTA) as its intracellular domain. This receptor harbors a myc-tag on its extracellular domain that can be visualized by immunostaining (Addgene *#*79128). A second virion expresses both the mCherry reporter and BFP under control of the Tetracycline responsive element (TRE-3G) promoter and the constitutive PGK promoter respectively. The third virion expresses surface GFP ligand under transcriptional control of the TRE-3G promoter. All cell lines were sorted for expression of the trans-genes. All cell cultures were maintained in an incubator at 37% humidity with 5% CO2. For viral transduction, cells were plated in 6-well dishes to achieve approximately 10% confluence at the time of infection. For lentiviral transduction, 10–100 ml of each virus supernatant was added directly to cells, with 1 *µ*l of polybrene (Millipore Sigma) also added to increase infection efficiency. Viral media was replaced with normal growth media 48 hours post-infection. Cells were sorted for co-expression of each component of the pathways via fluorescence-activated cell sorting (FACS) on a FacsAria2 (Beckton-Dickinson) and by staining for the appropriate myc-tag with fluorescence-tagged antibody where needed. A bulk-sorted population consisting of fluorescence-positive cells was established for “sender” cells. For single-cell clonal population establishment of transceivers, single cells were sorted by FACS into 96-well plates starting from populations of cells infected with lentiviral particles for the relevant expression constructs. After sorting, monoclonal population were expanded and screened for the activation of the GFP ligand after stimulation with anti-myc antibodies (Cell Signaling Technology) bound to an A/G plate (Thermo Scientific).

### Experiments

#### Signaling modulation assay

Senders and receivers L929 cells were co-cultured in a 1:1 ratio in DMEM for 24h. Then, cells were detached and analyzed by FACSAria2 (Beckton-Dickinson) as described in the Results. Details on experimental design. What were the stiffnesses of the reference, fibronectin, gelatin, Matrigel surfaces? What were the substrates for the reference, 0.2 kPa, 8,0 kPa, 64 kPA? Culture conditions? Cell densities? Concentrations of factors? Timing

#### Mechanism experiments

we measured cell shape and cell volume (via FACS for volume, via microscope imaging for area) (Fig. S3A-C); we measured GFP ligand accumulation via FACS and GFP ligand aggregates via image analysis of fluorescent microscopy images and correlate these with cell size (Fig. S3D-H); we estimated cell motility (Fig. S3I); we observed YAP localization (Fig. S3J); we measured signaling strength in presence of conditioned media from other confluences (Fig. S3K); and we measured nuclear marker fluorescent protein accumulation via FACS (Fig. S3L).

#### Propagation experiments

The L929 Sender and Transceiver cells, previously cultured in presence of tetracycline (100 ng/ml), were co-plated at ratio of 1:1000 in absence of tetracycline. Cell were imaged by automated inverted epifluorescence microscope (Keyence BZ-X710) at a magnification of 2X for the image acquisition of the whole well and at 20X for the image acquisition of the single spots. Gradients were generated by non-uniformly plating the cell suspension on one side of the well.

#### Growth curves

At day 0 cells were plated in 96 well plate at different densities, with and without drug treatments. Multiple replicates were plated to have one well for each day, until day 7. After imaging that was done to evaluate the extent of the propagation, cells were detached and resuspended in 100 *µ*l of PBS and counted three times by automatic cell counter (Countess II FL – life technologies).

### Image analysis

#### Processing of fluorescence images

To quantify the profiles of fluorescence intensity in microscope images, analysis was performed by Fiji-ImageJ. Background was subtracted and binary masks for fluorescent signals were generated to automatically segment propagation spots and quantify the area of fluorescence.

#### Cell shape analysis

We sought to quantify the change in projected cell shape. In the absence of a membrane marker, zoomed-in fields of view were selected at random from bright field images taken before FACS analysis (1:1 Sender:Receiver ratio, imaged at 24h of co-culture; see Figure S1C). Using Python, outlines of a subset of cells in the field of view were drawn manually and their area and perimeter were calculated. Both Senders and Receivers were counted and not distinguished. A previously described circularity index^84^ was also calculated as 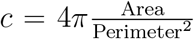. For a perfect circle, *c* = 1, and as the shape becomes less circular, *c* → 0.

#### Inference of cell motility

### Statistical analysis

#### Comparison of fluorescence measurements by FACS

Due to the hundreds to thousands of measurements per sample generated by flow cytometry, quantitative comparisons of the distribution of fluorescence values between samples can be highly sensitive to minor technical variations. To quantify the signaling state of Receivers using their FACS-measured mCherry fluorescence, fluorescence samples were compared to reference samples cultured with and without Sender cells (“ON” and “OFF” signaling states, respectively), both plated at 1x density. Empirical reference distributions were inferred by regularizing and binning the reference samples. Each distribution describes the statistical likelihood of observing a given fluorescence value in that condition. Then, for each data sample **x** = {*x*_*i*_}, a likelihood ratio LR was computed:

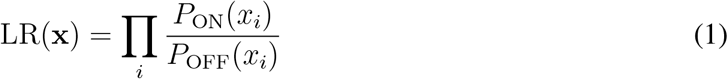

Here, *P*_ON_(*x*_*i*_) is the likelihood of observing the fluorescence value *x*_*i*_ if were drawn from the ON reference distribution (likewise for *P*_OFF_(*x*_*i*_)). In practice, the log-likelihood ratio LLR(**x**) = log_10_ LR(**x**) was calculated. Samples more likely to be OFF than ON (LR(**x**) < 1, or LLR(**x**) < 0) despite the presence of Senders were considered to have statistically significant suppression of signaling.

A perturbation was considered inhibitory (labeled **) if its observed fluorescence values **F** at 24 hrs were more likely to arise from the OFF condition than the ON condition, as measured by the parameter-free likelihood ratio 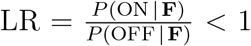, or log_10_ LR < 0 (Figure **1**). This procedure was found to be robust to the number of histogram bins (Figure S2A).

#### Estimation of population growth parameters

To model the dynamics of cell density in culture, we use the logistic equation, a differential equation that describes how population density *ρ*(*t*) evolves over time.

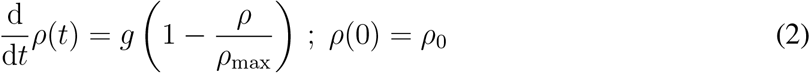

At low densities (0 < *ρ* << *ρ*_max_), there is exponential growth at the intrinsic growth rate *g*. As density increases further, however, population density saturates as it asymptotically approaches the carrying capacity *ρ*_max_. We then applied maximum likelihood estimation (MLE) to infer parameters of the logistic equation from the cell density measurements from growth curve experiments (see Methods:). All density counts were assumed to be normally distributed about the curve with constant standard deviation *σ* (homoscedastic). First, the parameters *g, ρ*_max_, and a were estimated using data from untreated co-cultures starting from three different starting densities. We then studied the effects of pharmacologic treatments by estimating *g* and *σ* in the presence of ROCK-i and FGF2. The carrying capacity *ρ*_max_ was held constant in order to alleviate degeneracies during fitting. Specifically, the logistic equation becomes underdetermined when the data has near-constant population dynamics. Due to measurement noise and low time-resolution, such data can correspond equally well to either very slow growth or very low carrying capacity, which obscure the true values of *ρ*_max_ and *g*, respectively. To resolve this degeneracy, carrying capacity (*ρ*_max_) was assumed to be constant under pharmacologic perturbation. Although these two possibilities suggest different regulatory influences on cell behavior, our model of signaling only takes into account the density itself and is agnostic to any underlying processes regulating it. Thus, either case will produce the same predicted signaling behavior.

## Mathematical modeling

### Dynamical equations for Receiver and Transceiver signaling

Following a dimensional analysis procedure (see Supplementary text), we model the dynamical response of a Transceiver cell using a system of delay differential equations.

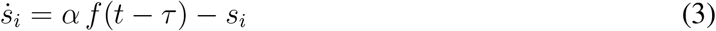

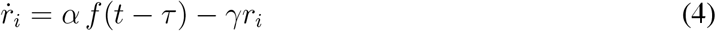

Here, for a given cell *i, s*_*i*_ and *r*_*i*_ denote the amounts of ligand (“signal”) and reporter, respectively. The maximum protein production rate *α* is equal for both species after non-dimensionalization. Protein degradation time-scales for ligand and reporter are *γ*_*s*_ and *γ*_*r*_, and the time-scale of reporter kinetics relative to ligand kinetics is represented by, and a time delay *τ* represents the time for protein manufacture and trafficking. Activation is taken to be a non-linear Hill-like function.

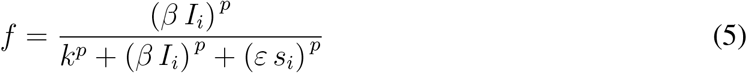

Here, *k* and *p* are the threshold and Hill coefficient (ultrasensitivity) of activation. A Hill-like activation function was chosen because this form can be derived from the application of mass-action kinetics to a cooperative transcription factor assembly, such as the dimerization of tTA-VP16. Given a cell *i* and its neighbors *j* ∈]*i*[, the total amount of input ligand is *I*_*i*_ = ∑_*j*_ *w*_*ij*_ *s*_*j*_, where *W* = (*w*_*ij*_) is the cell-cell contact matrix described below. The parameter *ε* is the strength of *cis*-inhibitory interactions by which the ligand inhibits synNotch activation in the same cell, indirectly inhibiting its own production. As the cell density *ρ* increases, the parameter *β* = exp (−*m*(*ρ* − 1)) dampens the effective amount of input. Table 1 shows the parameter values used for all simulations shown.

### The cell-cell contact matrix

The matrix *W* is a right-stochastic (Markov) matrix describing the amount of contact between cells *i* and *j*, where *i* is sensing ligand on the surface of *j*. Because of the Markov property ∑_*j*_ *w*_*ij*_ = **1**. Thus, *I*_*i*_ is a weighted mean of the ligand expressed by neighbors. Weights for each cell-cell pair were calculated using a Gaussian kernel centered on the signal-receiving cell, truncating the kernel at a maximum contact radius *r*_int_, and normalizing to satisfy the Markov property. Figure S5A shows an example of these weights.

### Integration of delay differential equations

Equations 3 were simulated using a fixed time-step forward-Euler integration scheme. Time delays were resolved using the method of steps. Initial conditions for signal expression were drawn from a random distribution.

### Cell density and length-scale in the model

In our model, we define the area of each hexagonal cell to be equal to the average area at the given density. For instance, at a density of 1250 cells/mm^2^ the cell area is 1250^−1^ mm^2^ = 800*µ*m^2^. The side length of a hexagon is then calculated from the area and used to infer linear distances on the plane of the lattice (propagation distances, scale bars, etc.). To circumvent computational complexity associated with re-meshing after each cell division, density changes are modeled by re-scaling cell sizes using the area relationship above, preserving the structure of the lattice. This choice may under-estimate propagation velocity due to the contribution of cell division to signal diffusion.

